# Efficient epistasis inference via higher-order covariance matrix factorization

**DOI:** 10.1101/2024.10.14.618287

**Authors:** Kai S. Shimagaki, John P. Barton

## Abstract

Epistasis can profoundly influence evolutionary dynamics. Temporal genetic data, consisting of sequences sampled repeatedly from a population over time, provides a unique resource to understand how epistasis shapes evolution. However, detecting epistatic interactions from sequence data is technically challenging. Existing methods for identifying epistasis are computationally demanding, limiting their applicability to real-world data. Here, we present a novel computational method for inferring epistasis that significantly reduces computational costs without sacrificing accuracy. We validated our approach in simulations and applied it to study HIV-1 evolution over multiple years in a data set of 16 individuals. There we observed a strong excess of negative epistatic interactions between beneficial mutations, especially mutations involved in immune escape. Our method is general and could be used to characterize epistasis in other large data sets.

## Introduction

Epistasis is common in nature and plays an important role in evolution^1,2^. In the presence of epistasis, the fitness effects of mutations are contingent on the genetic background in which they appear, making the relationship between sequence and function complex^3–5^. More accurate estimates of epistasis could improve our ability to predict evolution, both at the level of genetic sequences and phenotypes^6–8^.

Enormous amounts of time-resolved sequence data have been generated in recent years, opening up the possibility of inferring epistasis from observations of evolution. Naively, we anticipate that sets of mutations that improve fitness will be found together in the same genetic sequence more often than expected by chance, while sets of deleterious mutations will be observed less frequently. However, phenomena such as genetic hitchhiking^9^ and clonal interference^10^ can also generate correlations between mutations that are unrelated to function. At present, a few methods exist to estimate pairwise epistatic interactions from temporal data, but computational constraints limit their applicability to small numbers of loci^11–13^.

Here we propose an efficient method for inferring epistatic fitness that extends and vastly improves the computational efficiency of an approach developed by Sohail and collaborators^13^. With this new approach, the required memory and computational complexity scale only quadratically with the number of loci. These improvements are due to an efficient higher-order covariance matrix factorization (HCMF) method, which allows us to analyze much larger data sets than in previous analyses.^1–10^

After validating our method in simulations, we apply it to study epistasis in within-host human immunodeficiency virus (HIV)-1 evolution in a cohort of 16 individuals. Several past studies have highlighted the role of epistasis in viral evolution. Early experimental work found evidence for both synergistic^14^ and negative^15^ epistasis in different viruses. Epistasis has been observed in influenza and in SARS-CoV-2, especially in the context of immune evasion^16–21^. In HIV-1, epistasis has been observed between mutations involved in drug resistance^22–24^ and immune escape^25,26^. Here we found a consistent pattern of negative epistasis in HIV-1, with an interaction strength that typically scales along with the fitness effects of the individual mutations. Overall, our HCMF method enables the estimation of epistasis in large data sets, and our analysis contributes to the quantification of epistasis in viral evolution.

### Epistasis inference framework

As in related work^13,27^, our modeling framework is based on the Wright-Fisher (WF) model^28–30^. We write the number of haploid individuals with genotype *α* at time *t* in a population as *n*_*α*_(*t*). Given a total population size of *N*, we write the genotype frequency as *z*_*α*_(*t*) = *n*_*α*_(*t*)*/N*. The state of the population is described by frequencies ***z*** = (*z*_1_, *z*_2_, …, *z*_*M*_) = (*z*_*α*_(*t*))_*α*_ for each of the *M* possible genotypes. Its evolution is influenced by several factors, including mutation, recombination, and the fitness of each genotype. To model the fitness effects of individual mutations and pairwise epistasis, we assume a fitness function

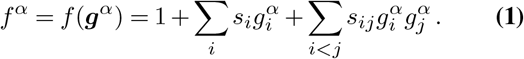

In the expression above, each locus *i* has a corresponding selection coefficient *s*_*i*_ that quantifies the fitness effect of the mutant allele at that locus and epistatic interactions *s*_*ij*_ with mutant alleles at all other loci *j*. For simplicity, we’ve used a binary model where each allele is either wild-type (WT) or mutant, but this can easily be extended to realistic sequence models (see Methods). Following this binary assumption, there are *M* = 2^*L*^ possible genotypes for sequences with *L* loci. The 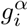 above are indicator functions, with a value equal to one if genotype *α* has a mutant allele at locus *i* and zero otherwise. Ultimately, our goal will be to infer the underlying fitness parameters 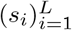 and (*s*_*ij*_)_*i<j*_ from temporal genetic data.

Under the WF model, the probability of obtaining a certain distribution of genotype frequencies in the next generation ***z***(*t* + 1), given the current distribution ***z***(*t*), is multinomial:

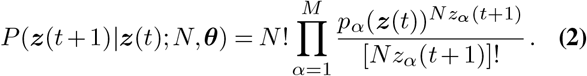

We use ***θ*** as a shorthand for all evolutionary parameters, including parameters describing selection and rates of mutation and recombination (Methods).

While inferring fitness parameters directly from (2) is challenging, when generation-to-generation changes in genotype frequencies are small, we can apply the simplified diffusion approximation of the WF model^31,32^. Through the diffusion approximation, we can obtain an analytically tractable expression for the probability of an evolutionary trajectory 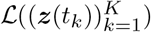, which we refer to as the path likelihood^13,27^ (Methods). This allows us to compute the fitness parameters (including individual selection coefficients *s*_*i*_ and pairwise epistatic interactions *s*_*ij*_) that best fit a data set of sequences collected over time.

To express the results, it’s useful to define a new vector ***s*** = (*s*_1_, *s*_2_, …, *s*_*L*_, *s*_1,2_, *s*_1,3_, …, *s*_*L−*1,*L*_) = (*s*_*e*_)_*e*_. This vector combines both selection coefficients for individual mutations and pairwise epistatic interactions, with a generalized index *e* that runs over both single loci (1, 2, …, *L*) and pairs of loci ((1, 2), (1, 3), …, (*L −* 1, *L*)). Similarly, we can define a mutant allele frequency vector ***x*** = (*x*_1_, …, *x*_*L*_, *x*_1,2_, …, *x*_*L−*1,*L*_), where

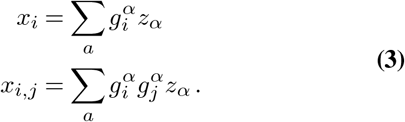

Higher order mutant allele frequencies (e.g., *x*_*i,j,k*_) are defined similarly.

The fitness parameters *ŝ* that maximize the path likelihood, together with a Gaussian prior distribution (or equivalently, ridge regression penalty) for the selection coefficients and epistatic interactions, are then given by^13,27^

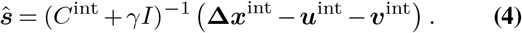

In this expression, the observed net allele frequency change over the observation period is

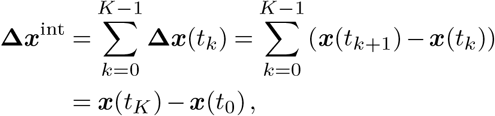

while ***u***^int^ and ***v***^int^ represent expected cumulative frequency changes due to mutation and recombination, respectively. Explicit derivations and definitions for these terms are given in Methods. *C*^int^ is the allele frequency covariance matrix integrated over time, and *γ* quantifies the width of the prior distribution for the selection coefficients and epistatic interactions.

Although (4) is complicated, it can be interpreted intuitively. Essentially, (4) states that allele frequency change that is *not* explained by the forces of mutation or recombination is evidence of selection. The sign and magnitude of inferred selection depend on the net change in allele frequencies (how much, quantified by Δ***x***^int^, and how fast, quantified by the diagonal part of *C*^int^) as well as the effects of genetic background (quantified by the off-diagonal terms of *C*^int^).

## Results

### Factorization of higher-order integrated covariance matrix and efficient inference framework

While (4) provides a powerful expression to simultaneously estimate the fitness effects of mutations and pairwise epistasis from temporal genetic data, it faces a serious computational limitation. The covariance matrix *C*^int^ is *D* = *qL*(*qL*+1)*/*2-dimensional, where *q* is the number of alleles at each locus. The computational complexity of inverting the covariance matrix thus scales as *𝒪* ((*qL*)^6^), with memory costs related to the storage of covariance matrix entries scaling as *𝒪* ((*qL*)^4^). For data sets with hundreds or thousands of loci, simply storing the covariance matrix in memory becomes challenging.

We developed an efficient and generic method to resolve the major computational bottleneck hindering the application of this approach to larger data sets. The key idea of our approach is to exploit the regular structure of the covariance matrix, allowing us to factorize the matrix and perform calculations in a lower-dimensional space without any loss of information. Specifically, we can write the integrated covariance matrix in terms of a rectangular matrix with dimensions *D* × *d*, which depends on the number of unique sequences in the data set. Writing the number of unique sequences in the data set at time *t*_*k*_ as *d*_*k*_, we have 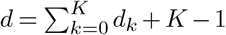. As we will show below, *d* may be multiple orders of magnitude smaller than *D* for relevant data sets of interest, allowing this factorization to dramatically speed up analyses.

In our analysis, we integrate the covariance matrix over the evolution using linear interpolation between sampling times, which mitigates periods of sparse sampling^13,27^. The integrated covariance matrix with linear interpolation can be factorized as follows (see Methods for details):

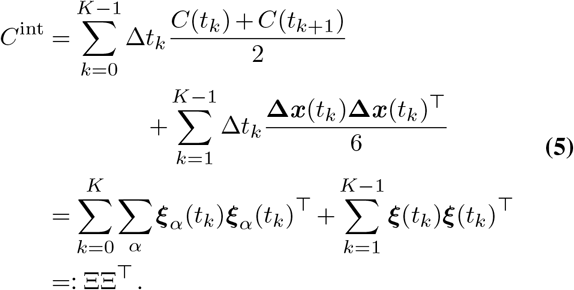

The ***ξ*** vectors are defined as

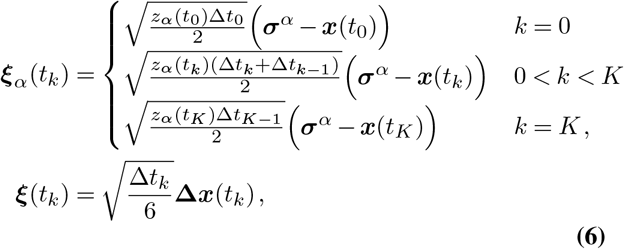

where Δ*t*_*k*_ = *t*_*k*+1_ *− t*_*k*_, *z*_*α*_(*t*_*k*_) is the frequency of genotype *α* in the data at time *t*_*k*_, and ***σ***^*α*^ is a *D*-dimensional vector with entries

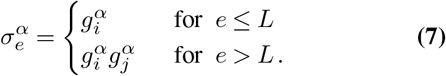

More complex covariance interpolation using spline curves^33^ can also be expressed in a form similar to (5).

Using the factorized Ξ matrix (5), we can rewrite the equation for the estimated selection coefficients and epistatic interactions (4) as

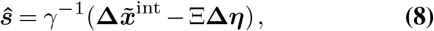

With

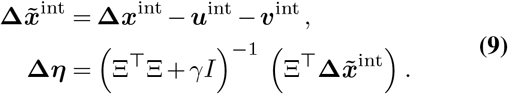

Critically, computing (8) is far less computationally intensive than (4) when *D* ≫ *d*, as the matrix to be inverted in (8) is only *d* × *d*. In total, the computational complexities of calculations in (8) are: matrix-vector products of ΞΔ***η*** and 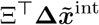 take *𝒪*(*dD*); matrix-matrix product of Ξ^⊤^Ξ requires *𝒪*(*d*^2^*D*); solving the equation for Δ***η*** without directly solving its inverse is smaller than *𝒪*(*d*^2+*ω*^), with *ω* a small positive number 0 *< ω* ≤ 1, depending on linear optimization solvers.

This substantial computational reduction was achieved by implicitly computing the integrated covariance matrix without ever storing the covariance matrix itself. Therefore, our epistasis inference scheme is more efficient and scalable as the computational complexity scales only linearly with *D* (and thus quadratically with *L*). In comparison, even naive selection inference without epistasis scales as *D*^3^. The expression for the selection coefficients in (8) uses no approximations. Thus, its solution is exact in the diffusion limit^13^.

For simplicity, we initially assumed the same regularization *γ* for selection and epistasis. However, we have also generalized our approach so that the regularization values *γ*_*e*_ can differ and implemented this in our code (Methods). While our analysis considers only pairwise epistatic interactions, one could further extend the fitness function to consider even higher-order interactions. For *p*-way epistatic interactions, the computational complexity would become *𝒪*(*dD*) with 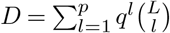.

### HCMF substantially reduces computational costs

To assess the efficiency of HCMF, we simulated population evolution under the WF model using different numbers of loci, ranging from *L* = 50 to 1600. We used a constant population size of *N* = 10^3^, a mutation rate of *µ* = 10^*−*3^, and a recombination rate of *r* = 10^*−*4^ per site per generation. Our simulations ranged over 2000 generations, with virtual samples collected for inference every 10 generations. We used a fitness landscape in which 25% of mutations were beneficial (*s*_*i*_ = 0.03), 25% were deleterious (*s*_*i*_ = −0.03), and 50% were neutral (*s*_*i*_ = 0). Similarly, 25% of all pairs of sites were randomly selected to have positive/negative epistatic interactions (*s*_*ij*_ = 0.03 or *−*0.03, respectively), with the remaining 50% of the possible epistatic interactions set to zero. To ensure sufficient sampling to measure typical results, we performed 500 simulations for each condition.

Since the size of the covariance matrix increases quadratically with sequence length, the required memory size of the naive approach increases as *𝒪*(*L*^4^). However, memory requirements only scale as *𝒪*(*L*^2^) for the HCMF method (**Fig. 1a**). HCMF also dramatically reduces the run time of the inference, scaling as *𝒪*(*L*^2^) compared to *𝒪*(*L*^6^) for the naive approach (**Fig. 1b**). For example, for *L* = 400, HCMF is 10^4^ times faster than the naive approach. This computational advantage should further increase for larger sequence lengths. As noted above, since the HCMF approach involves no approximations, the selection coefficients and epistatic interactions inferred by this approach match the ones from the naive method exactly within machine precision.

**Fig. 1.**
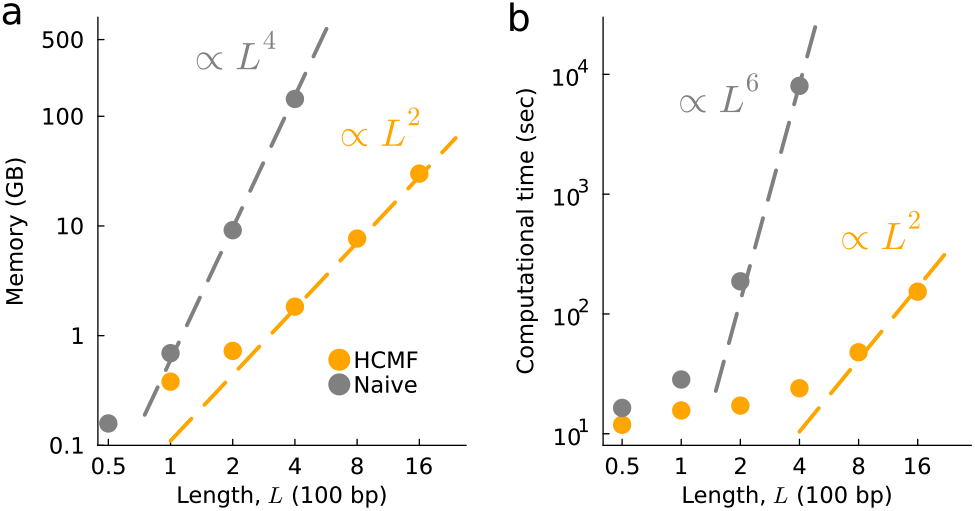
HCMF substantially reduces the required memory size and computational time. **a**, Required memory size versus number of loci *L* (measured in 100 bp). The required memory size of the naive method scales as *𝒪*(*L*^4^), while our method reduces it to *𝒪*(*L*^2^). For the naive method, we did not consider *L >* 400 due to computational constraints. **b**, Required computational time (in seconds) versus number of loci. As anticipated, the computational time of the naive method and HCMF scale by *𝒪*(*L*^6^) and *𝒪*(*L*^2^), respectively.

### Importance of higher-order covariance information for inferring epistasis

One of the main barriers to inferring epistatic interactions via (4) is computing and inverting the integrated covariance matrix. The HCMF approach we have developed offers one solution to this problem. However, one could also simplify (4) by neglecting the off-diagonal terms of the covariance matrix. This greatly reduces the computational burden of the problem, but neglects important information about linkage disequilibrium that could inform the inference of selection coefficients and epistatic interactions. We refer to this approximation as the independent model (analogous to the single locus model of ref.^27^).

We performed additional simulations to compare the accuracy of the HCMF method, which includes higher-order covariance information, and the independent model, which does not. These simulations were performed with the same parameters as in the previous section, using *L* = 50 loci and sparser epistatic interactions. Here we chose a random set of *L/*2 = 25 pairs of sites to have positive epistatic interactions (*s*_*ij*_ = 0.03), *L/*2 pairs with negative interactions (*s*_*ij*_ = *−*0.03), and set the remaining epistatic interactions to zero.

The inferred epistatic interactions using the full model with HCMF are much closer to the true ones than those inferred with the independent model (**Fig. 2a-b**). For the full model, the distribution of inferred positive/neutral/negative epistatic interactions is roughly normal, with peaks that can easily be distinguished from one another. In contrast, the epistatic coefficients inferred using the independent model are distributed much more broadly and irregularly. While positive and negative interactions can, on average, still be distinguished from one another using the independent model, it is more difficult to do so. To quantify this difference, we computed the receiver operating characteristic (ROC) curve and area under the curve (AUC) for identifying positive (**Fig. 2c**) and negative (**Fig. 2d**) epistatic interactions using the full and independent models. Thus, by all metrics we find that the inclusion of higher-order covariance information improves the ability to identify epistatic interactions from data.

**Fig. 2.**
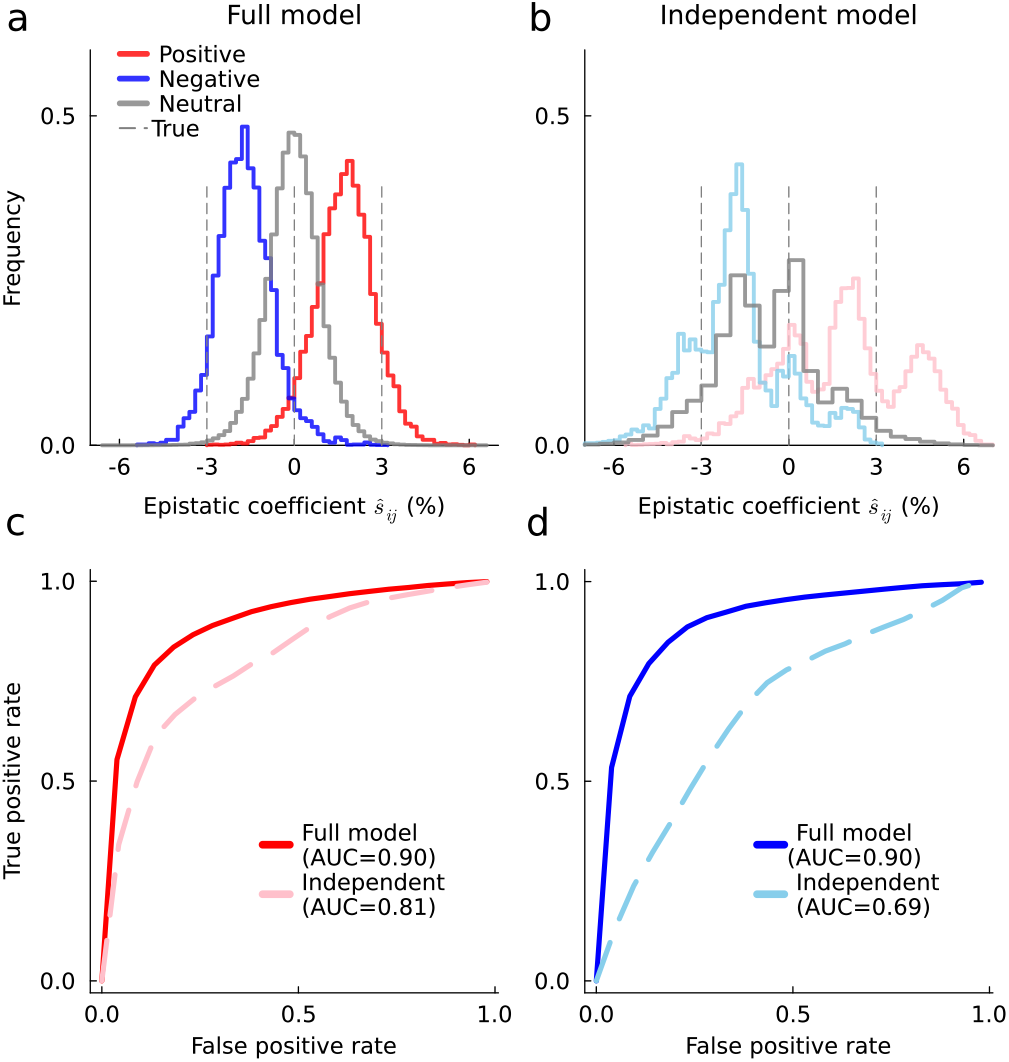
Higher-order covariance information improves the inference of epistasis. Distribution of inferred epistasis using the full model inferred via HCMF (**a**), which includes higher-order covariance information, and using the independent model (**b**), in which off-diagonal terms of the covariance matrix are set to zero. **c**, Receiver operating characteristic (ROC) curve for identifying positive epistasis. The area under the curve (AUC) value is 0.90 for the full model, while it drops to 0.81 for the independent model. **d**, Analogous ROC and AUC values for identifying negative epistasis. The AUC values are 0.90 and 0.69 for full and independent models, respectively.

### Modeling epistasis improves the inference of selection coefficients

An alternative approach to reducing the computational costs of (4) is to use a simpler fitness landscape, such as a purely additive one with all *s*_*ij*_ = 0. This assumption may be especially appropriate for analyzing highly similar sequences. However, in models with substantial, strong epistatic interactions, omitting epistasis could skew fitness estimates (**Fig. 3a-b**). In these conditions, inference using models with epistasis yields more accurate estimates of individual selection coefficients and improves the detection of beneficial and deleterious mutations (**Fig. 3c-d**).

**Fig. 3.**
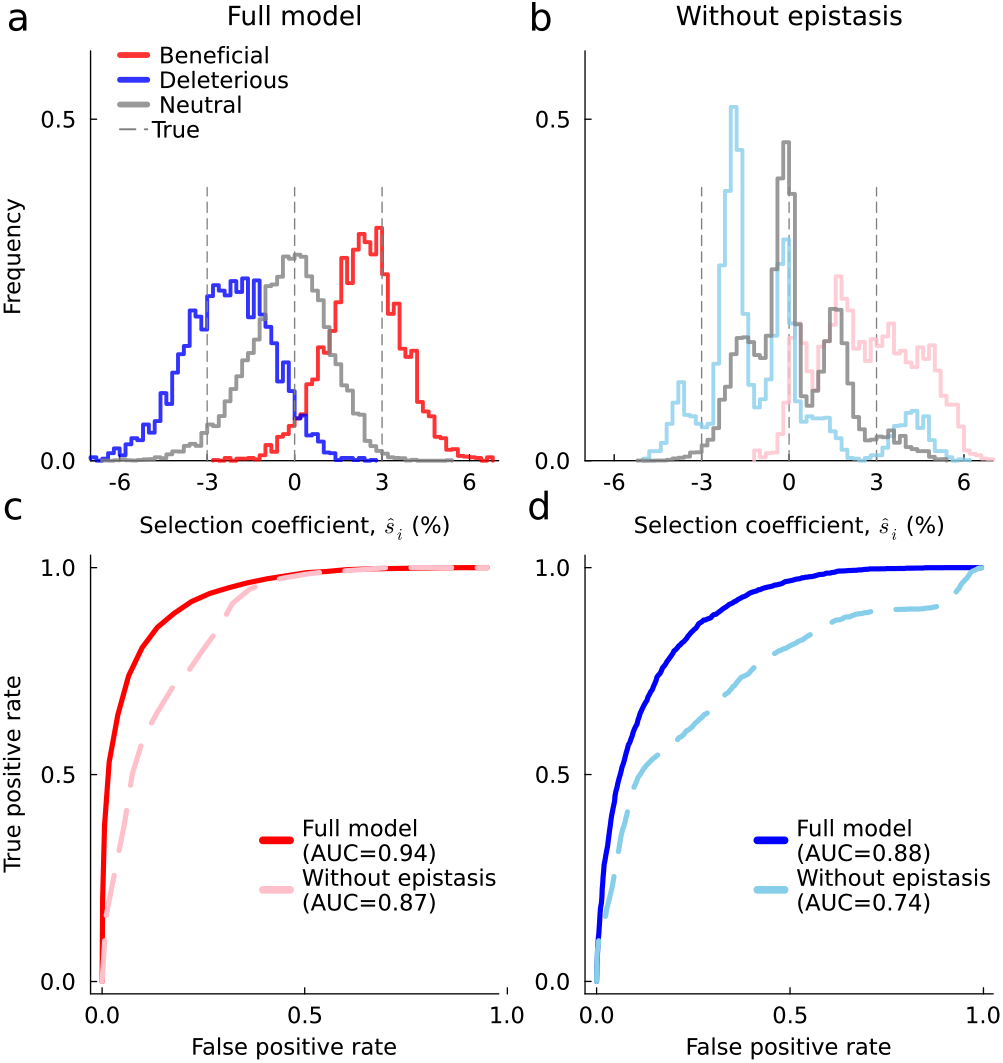
Modeling epistasis improves the inference of additive selection coefficients. Distribution of selection coefficients inferred with the full model via HCMF (**a**) and a simpler model with no epistatic interactions (**b**). When epistasis is present, including it in the model also improves estimates of selection coefficients. **c**, ROC curves and their AUC values for identifying positive selection coefficients. The AUC value of the full model is 0.94, while the AUC value of the model without epistasis drops to 0.87. **d**, Analogous ROC and AUC values for identifying deleterious selection coefficients. The AUC values are 0.88 and 0.74 for the full model and the model without epistasis, respectively. Simulation parameters are the same as in **Fig. 2**.

### Joint inference from multiple replicates

Some evolution experiments, such as deep mutational scanning studies^34,35^, have multiple independent replicates collected under the same conditions. Using our approach, we can estimate selection coefficients and epistatic interactions that best explain the data across all replicates, as shown in prior work^13,36,37^. To demonstrate this phenomenon in a challenging setting for inference, we increased the density of epistatic interactions to 50%, with half set to be positive (*s*_*ij*_ = 3%) and half negative (*s*_*ij*_ = *−*3%). In this case, epistatic effects dominate the fitness function. With a single replicate, the AUC values were 0.82 (0.81) for identifying beneficial (deleterious) selection coefficients and 0.74 (0.72) for positive (negative) epistasis. Combining data from two replicates raised the AUC to 0.93 (0.89) for selection coefficients and 0.86 (0.85) for epistasis, with further improvements as the number of replicates increases (**Supplementary Fig. 1**).

### Epistasis in intrahost HIV-1 evolution

As a practical application of our approach, we studied within-host HIV-1 evolution in 16 individuals who were not treated with antiretroviral drugs during the sampling time. This data set included individuals enrolled in the CHAVI 001 and CAPRISA 002 studies in the United States, Malawi, and South Africa^38,39^. Each individual was identified shortly after HIV-1 infection, and the viral population within each individual was sampled frequently for several months to years afterward. For most individuals, the 3^*′*^ and 5^*′*^ halves of the HIV-1 genome were sequenced separately using single genome amplification methods, preserving information about linkage disequilibrium between mutations even at long distances. For two individuals, denoted CH505 and CAP256, only the HIV-1 surface protein Env was sequenced. Most data sets consisted of around 50-100 HIV-1 sequences in total for each sequencing region, collected over 5-8 time points, with several hundred polymorphic loci (see **Supplementary Table 1**). However, the viral population was also sequenced more deeply in a few individuals, featuring as many as 1205 HIV-1 sequences collected at 31 time points over roughly 5 years.

Using this data, we inferred selection coefficients and epistatic interactions between HIV-1 mutations for each individual and sampling region. We used prior estimates to set the mutation^40^ and effective recombination rates^41^ in (8). By convention, we set the selection coefficients and epistatic interactions for the transmitted/founder (TF) sequence, the natural analog of WT, to zero (Methods). Thus, fitness effects are expressed relative to the strain of the virus that originally infected each individual. In general, the ability to transform the model parameters (i.e., selection coefficients and epistatic interactions) without affecting the dynamics of the model is referred to as a gauge freedom. Choosing a specific convention for the parameters is important for comparing fitness effects in different contexts and for improving the interpretability of the model^37,42–44^.

Here we focused specifically on epistatic interactions between nearby sites (separated by <50 bp), with distant epistatic interactions suppressed by strong regularization. There were two reasons for our focus on short-range interactions. First, due to the high effective recombination rate in HIV-1, the size of the sequencing region, and some large time gaps between samples, the expected change in correlations between mutant alleles due to recombination may violate the mathematical assumptions of the diffusion approximation, biasing our inferences for these sites. Second, short-range epistatic interactions may be of particular biological interest in HIV-1 evolution.

The accumulation of mutations within cytotoxic T lymphocyte (CTL) epitopes – linear peptides roughly 10 amino acids in length that are recognized by cytotoxic T cells – allows mutant viruses to escape from the immune system. Past work has shown that T cells are especially important in controlling HIV-1 replication^45^, and that the virus faces significant selective pressure to escape from CTLs ^27,39,45–47^. However, because the recognition of CTL epitopes is highly specific, even one nonsynonymous mutation within the epitope can be sufficient to confer escape^48–50^. We anticipate that this phenomenon could lead to negative epistasis between CTL escape mutations, as the fitness benefit of multiple mutations within the epitope should be lower than expected based on the beneficial effect of each individual escape mutation.

While most of the epistatic interactions we inferred were very close to zero, a few were significantly negative (**Fig. 4**). We observed a general trend where negative epistatic interactions were more common between beneficial mutations, especially CTL escape mutations (**Fig. 4**). This pattern of negative epistasis between beneficial mutations, including CTL escape mutations, was robustly observed across all individuals and sequencing regions that we studied (**Supplementary Fig. 2**).

**Fig. 4.**
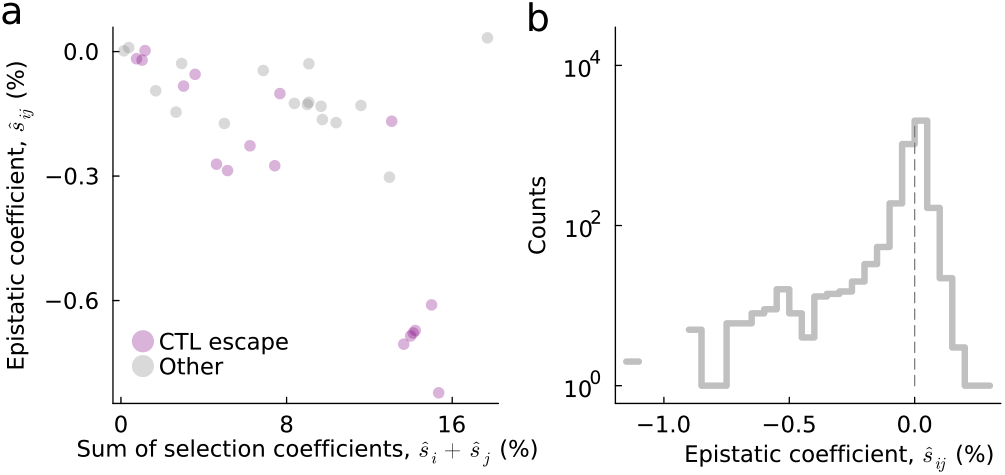
Predominance of negative epistasis between beneficial HIV-1 mutations. **a**, Comparison of inferred epistatic interactions, *ŝ*_*ij*_, and the corresponding sum of individual selection coefficients, *ŝ*_*i*_ + *ŝ*_*j*_, in a typical case (700010077-3; see **Supplementary Fig. 2** for all individuals). Generically, we find negative correlations between inferred epistatic interactions and selection coefficients. CTL escape mutations are typically found to be both strongly beneficial and to have negative epistatic interactions with other escape mutations. **b**, Distribution of inferred epistatic interactions across all individuals. Most terms are near zero, but a few epistatic interactions are significantly negative.

### Consistency with prior estimates of selection in HIV-1

Past work has studied HIV-1 evolution in part of this data set with different modeling choices, including a model with purely additive selection^27^ and one that includes specific terms for CTL escape^51^. Neither of these models includes pairwise epistatic interactions. Thus, we compared the selection coefficients inferred in our analysis with those from previous models to understand how the inclusion of pairwise epistasis affects the interpretation of the fitness effects of individual mutations.

**Figure 5a** shows a typical example of the inferred selection coefficients, *ŝ*_*i*_, with and without the inclusion of epistasis. Overall, we find that the inferred selection coefficients are similar to those in past models (mean Pearson’s *R* = 0.94). In particular, all models find very strong selection for CTL escape mutations^27,51^. As in previous work, the great majority of inferred selection coefficients are very close to zero (**Fig. 5b**). However, the model without epistasis also features heavier tails in the distribution of inferred selection coefficients, with more mutations inferred to have either very beneficial or very deleterious individual effects.

**Fig. 5.**
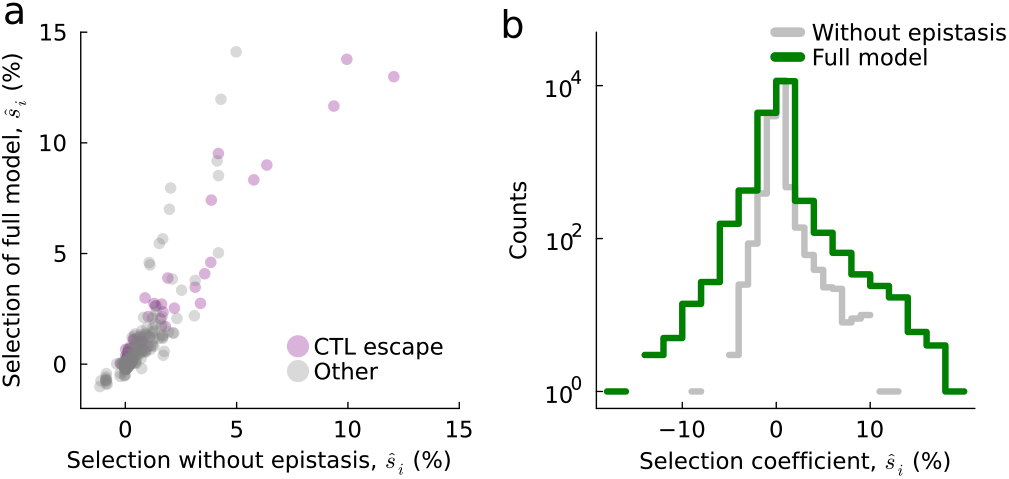
Consistency of inferred selection coefficients in models with and without epistasis. **a**, Comparison of inferred selection coefficients in models with and without epistasis in a typical case (700010077-3; see **Supplementary Fig. 3** for all individuals). While the exact values differ, there is excellent general agreement between the mutations that are inferred to strongly affect fitness and those that are inferred to be nearly neutral. **b**, Distribution of selection coefficients across all individuals. Both distributions are peaked near zero, but the tails of the distributions in the full model are longer.

## Discussion

Epistasis is prevalent in nature, and has been observed to influence viral evolution ^1,15,52–54^. However, inferring epistasis from data is technically and computationally challenging. Here, we developed a new approach for the path likelihood inference framework ^13,27,37,51,55^ that greatly reduces computational costs for many data sets of interest, especially for inferring epistasis. Our key innovation was the efficient factorization of the higher-order covariance matrix, which allows us to analytically estimate selection coefficients and epistatic interactions from data without ever explicitly computing the covariance matrix or its inverse. For this reason, we referred to our method as higher-order matrix factorization (HCMF). The HCMF approach is general and can be applied under different assumptions about the structure of the fitness landscape. HCMF does not introduce any new approximations, so it suffers no loss in accuracy compared to prior approaches.

After validating our approach in simulations, we applied HCMF to study HIV-1 evolution within 16 individuals. The fitness effects of mutations that we inferred were consistent with past computational results^27,36,47,51^ and with experimental findings. In particular, we found strong selection for mutations that allow the virus to escape from the host immune system, in agreement with a large body of experimental work and clinical observations ^25,39,45,46,56^.

In this HIV-1 data set, the distribution of epistatic interactions that we inferred was peaked near zero, but with a substantial tail of strong negative epistasis. Patterns of negative epistasis have also been observed in other viruses^15^. Negative epistasis was especially common between CTL escape mutations, consistent with the finding that single mutations within an epitope typically already disrupt T cell recognition^48–50^. We also observed negative epistasis between pairs of beneficial mutations more generally. This finding is consistent with more general studies that have observed decreasing effect sizes of beneficial mutations over time^6,57–59^.

Our approach to inferring epistasis from temporal data differs from some prior methods, which used statistical models to explain correlations in protein sequences collected from many individuals or species ^22,23,26,60–64^. These models treat sequence data as samples from a static, equilibrium distribution, and interpret correlations between mutations as possible evidence for epistasis. Only a handful of methods allow for the possibility that linkage disequilibrium may arise from an underlying phylogenetic structure to the sequence data^65,66^, or simply by chance. Nonetheless, these approaches have also been successful at tasks such as predicting the fitness effects of HIV-1 mutations in experiments^62,63^ and the dynamics of immune escape within individual patients^26^. In contrast to the present work, these models offer a “global” view of epistasis averaged across many related sequences.

The HCMF approach that we have developed is general. While our study focused on HIV-1, future work could be applied to other populations, including viruses like influenza and SARS-CoV-2 (ref.^55^), experimental evolution^37^, or bacteria^67,68^. As one example, recent studies have suggested that epistasis plays an important role in maintaining fitness among SARS-CoV-2 Spike mutations that escape from antibodies and control receptor binding^19,20^. More systematic studies could reveal the importance of epistasis in different aspects of SARS-CoV-2 evolution. More generally, a deeper understanding of epistasis may also improve our ability to understand and predict viral evolution.

## ACKNOWLEDGEMENTS

The work of K.S.S. and J.P.B. reported in this publication was supported by the National Institute of General Medical Sciences of the National Institutes of Health under Award Number R35GM138233.

## AUTHOR CONTRIBUTIONS

All authors contributed to research design, methods development, interpretation of results, and writing the paper. K.S.S. performed simulations and computational analyses. J.P.B. supervised the project.

## Supplementary Information

## Methods

### HIV sequences

We retrieved HIV-1 sequences for individuals in this study from the Los Alamos National Laboratory (LANL) HIV Sequence Database^69^ (see **Supplementary Table 1**). We processed the sequence data as described in ref.^70^ to minimize the influence of sampling noise. Processing steps included 1) removing the sequences with large deletions, 2) removing sites with high gap frequencies (indicating rare insertions or potential alignment errors), and 3) eliminating time points with <4 sequences or ones that were obtained >200 days after the prior sampling time. In addition, we imputed ambiguous nucleotides using the most common nucleotides in the data at the same site within that individual.

### Model

We model the evolution of a population of *N* individuals subject to mutation, recombination, natural selection, and genetic drift (finite population size), following the Wright-Fisher (WF) model^71–73^. We assume that all individuals are haploids. Each genotype ***g*** ∈ *A*^*L*^ is a sequence of *L* alleles, with the alleles considered to be categorical variables *𝒜* = {A, T, G, C, -} for DNA or *𝒜* = {A, C, …, W, Y, -} for amino acids, including a gap character to represent deletions. We write the fitness of each genotype as

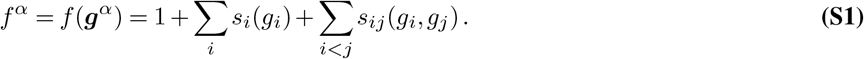

The *s*_*i*_(*a*) are additive fitness parameters acting on each site and allele individually, while the *s*_*ij*_(*a, b*) are epistatic interactions between pairs of alleles *a* and *b* at sites *i* and *j*. We assume the underlying fitness parameters are constant in time, with the population’s adaptation speed much faster than the rate of environmental changes. We define the probability of mutation from allele *a* to *b* as *µ*_*ab*_ per site per generation, which we assume is the same for all sites. Here we used a simple model of recombination, in which there is a probability *r* per site per generation for a recombination breakpoint to occur at that site. A recombinant sequence derived from two sequences ***g***^*α*^ and ***g***^*β*^ with recombination breakpoint *i* then has the form 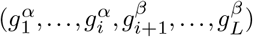. We assume that the partner sequence *β* is always chosen randomly from the population with a probability proportional to the frequency of that genotype.

Let ***z*** ∈ [0, 1]^*M*^ be the genotype frequency with the number of genotypes being *M*. The WF model under the fixed population size *N* is defined as the following multinomial process:

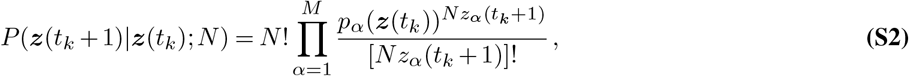

where *p*_*α*_ is given by

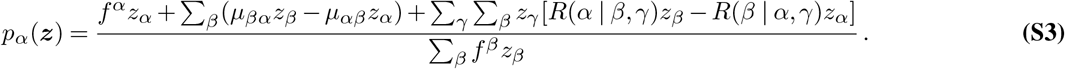

*R*(*α*|*β, γ*) is the probability that the recombination of genotypes *β* and *γ* results in a genotype *α*.

In simulations, we used *µ*_*ab*_ = *µ* = 10^*−*3^ and *r* = 10^*−*4^ per site per generation. For HIV-1 data analysis, we used mutation rates estimated from a longitudinal virus evolution study^74^, along with a constant recombination rate of *r* = 10^*−*5^ per site per generation, in line with past estimates of the effective recombination rate^75–79^. This choice for representing recombination in

HIV-1 is a simplification. In reality, HIV-1 recombination occurs in multiple steps: first, two different viruses must coinfect the same cell. Then, genetic material from each virus can be packaged together in the same virion. When such a virion infects a new cell, recombination can occur as the viral reverse transcriptase switches between templates. Thus, the effective HIV-1 recombination rate involves both coinfection and template switching probabilities. Recent work has also shown that the effective recombination rate can increase when viral load is higher, due to increased rates of coinfection^80^. Here we applied only the simple recombination model in which probabilities of coinfection and template switching are combined into a single effective recombination rate. Future work could relax this assumption and consider the effects of time-varying recombination rates due to fluctuations in viral load.

### Diffusion limit

The properties of multinomial processes lead to the following genotype average and covariance values:

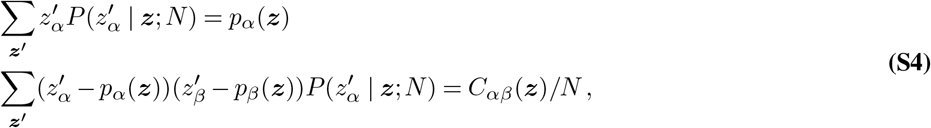

with

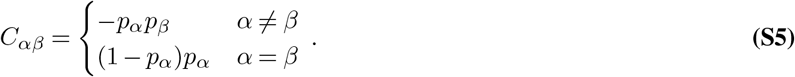

Assuming that the rates of mutation and recombination and the fitness effects of mutations are small (formally, *𝒪*(1*/N*)), changes in genotype frequencies are not abrupt, and we can employ the diffusion approximation^81^ to simplify the WF model. This results in the following Kimura’s diffusion equation (Fokker-Planck equation or Kolmogorov forward equation^82^):

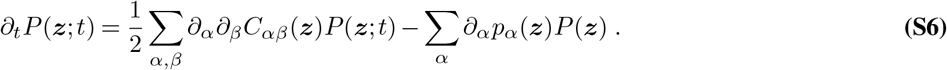

The above equation leads to the Gaussian process with average drift and diffusion matrix (Eq. (S4)). To be more explicit, assuming 1 ≪ *N* and collecting the only *𝒪*(1*/N*) terms in the stochastic process, we get the following tractable expression

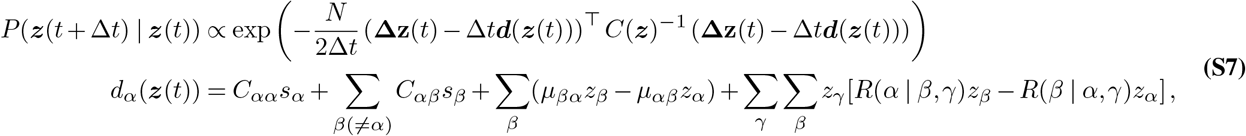

Here the covariance is also taking only the *𝒪*(1*/N*) terms and scaled by *N*, therefore, *C*_*αα*_ = *z*_*α*_(1 *− z*_*α*_) and *C*_*αβ*_ = *−z*_*α*_*z*_*β*_. In the main text, we gave the optimal selection coefficients *ŝ*_*i*_ and epistatic interactions *ŝ*_*ij*_ maximizing a posterior distribution over the above diffusion processes (Eq. (4)).

So far, we have discussed the diffusion process in the genotype distribution space. To make the expressions more transparent, we can project the genotype frequency dynamics onto the allele frequency space (Eq. (3)), which we describe below.

**Expected frequency change due to mutation**

Assuming the WF process, we can analytically estimate the expected frequency change due to mutation and recombination effects and integrate them over the generations. Define the indicator function 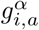 that gives one if genotype *α* has allele *a* at site *i*, otherwise zero, and *x*_*i,a*_ is the allele frequency obtained by 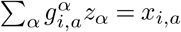.

Since the mutation rate is small, effectively, no more than one mutation occurs per generation for individual sequences.

Therefore, possible mutations between genotypes *α* and *β* are formally constrained by their distance such that 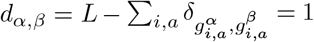, with *δ*_*x,y*_ being Kronecker’s delta. Therefore, the expected additive frequency change in allele *a* at site *i* due to mutation is:

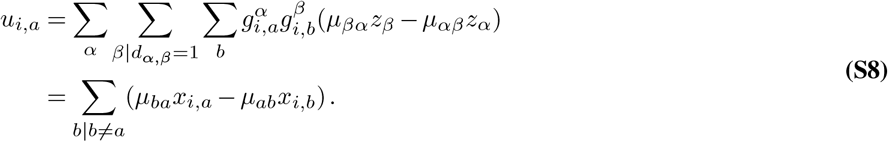

Similarly, for pairwise frequencies we obtain

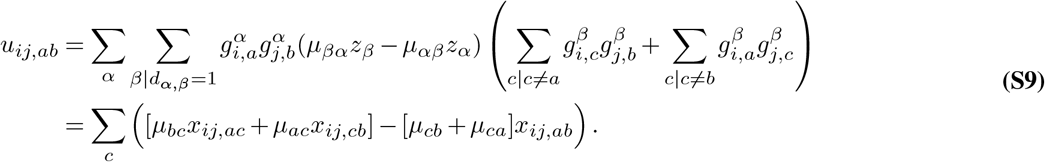

### Expected frequency change due to recombination

By symmetry, one can show that recombination has no effect on the expected change in individual allele frequencies^70^. However, correlations between mutations are naturally diluted by recombination. For pairwise frequencies, recombination decreases correlations between mutations until they become independent. One can show that the expected change in pairwise allele frequencies due to recombination is^83^

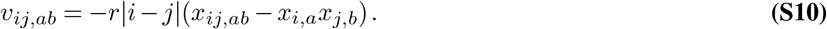

A detailed derivation is included in ref.^83^.

### Maximum *a posteriori* solution over the path

Leveraging the analytically tractable transition probability under the diffusion limit (Eq. (S7)), the maximum *a posteriori* estimate of the selection coefficients and epistatic interactions is^83^

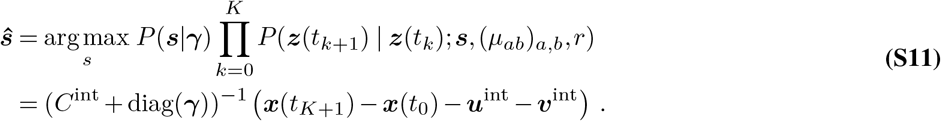

Here diag(***γ***) is a matrix with ***γ*** on the diagonal and zeros elsewhere. *P* (***s*** | ***γ***) represents a prior distribution for the selection and epistatic coefficients, given by the normal distribution

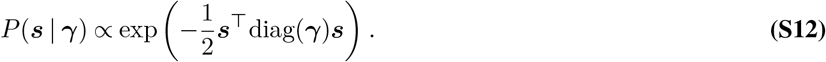

The expected net frequency change due to mutation and recombination, denoted by ***u***^int^ and ***v***^int^, are defined in the equation Eq. (S8), Eq. (S9) and Eq. (S10), respectively. Further reduction based on the HCMF method is explained in the main text. We provide the expression of the factorized covariance matrix in the following section.

### Representing the integrated covariance matrix with linear interpolation by a low-rank matrix

In this section, we show the expression of the integrated covariance matrix with piece-wise linear interpolation is given as the integration of covariance with a piece-wise constant interpolation and a sum of rank-one matrices.

By integrating them over the time *t*, we get the (time-) integrated covariance matrix. More specifically, we consider the following linear interpolation, such that

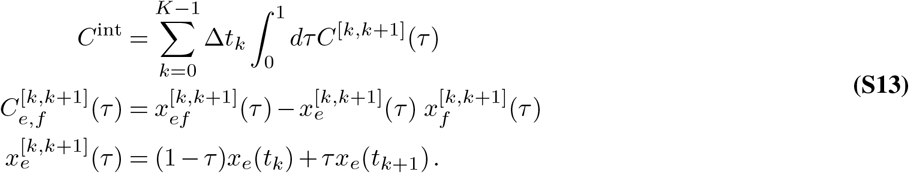

It is straightforward to check that the following expression is identical to the diagonal of the integrated covariance matrix with the piece-wise linear interpolation given in ref.^70^:

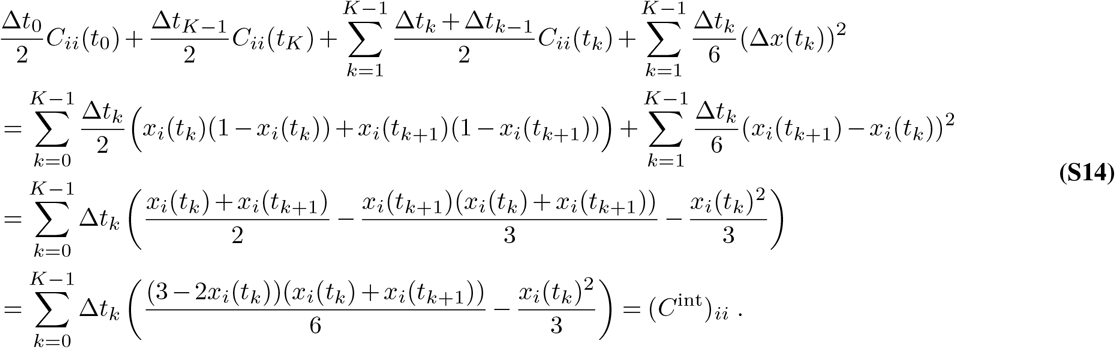

For the off-diagonal case, the pairwise-frequency term *x*_*ij*_(*t*_*k*_) is linear in time, and the result of the integral with the linear interpolation is the same as the integral with the piecewise-constant interpolation. Therefore, we explicitly write only the integrals that are non-linear in time:

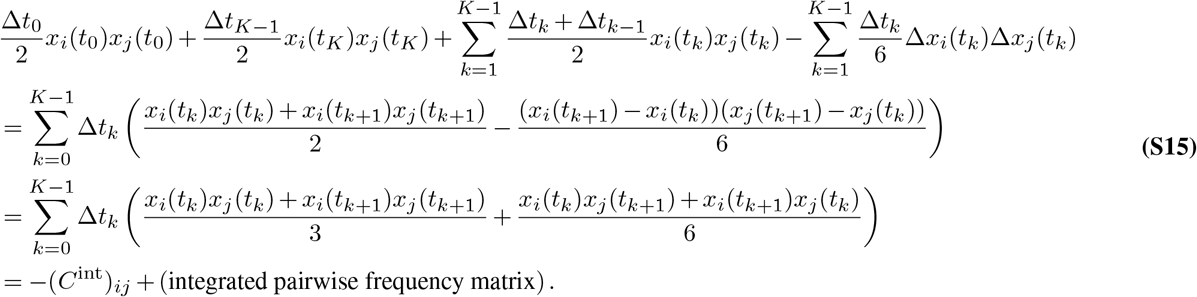

By summarizing these equations, we represent the integrated covariance matrix as:

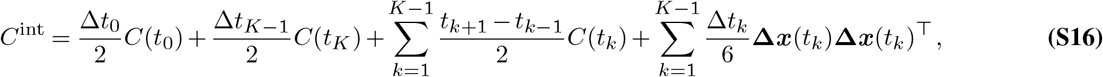

which we can readily factorize by a matrix Ξ such that *C*^int^ = ΞΞ^⊤^. The size of the matrix Ξ is *D* × *d*, where 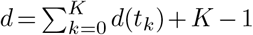 with *d*(*t*_*k*_) denoted as a rank of *C*(*t*_*k*_). In most of the evolutionary data, the size of the higher-order covariance matrix is much larger than the effective matrix rank size; hence, typically *d* ≪ *D*.

### Inferring fitness parameters from multiple replicate trajectories

In cases where multiple ensembles of trajectories evolve under similar conditions, it is natural to extend the path likelihood to multiple ensembles. Suppose there are *Q* replicates, let *q* be the index of the *q*th replicate, 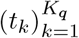 be a set of sampling time-steps for the *q*th replicate, and ***x***^*q*^(*t*_*k*_) be the set of single and pairwise frequencies for the *q*th replicate.

The maximum path likelihood solution using *Q* replicates can be expressed as^83^

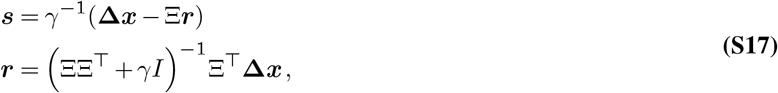

Where

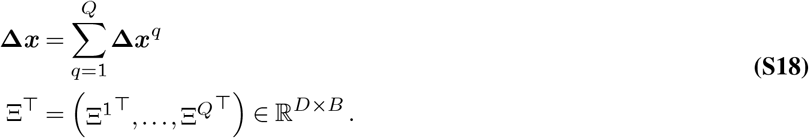

*B* is the total number of samples across replicates over the evolution, formally, 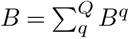, where *B*^*q*^ is the total number of samples of the *q*-th replicate over its evolution. Intuitively, the likelihood of multiple independent trajectories is equal to the product of each of their likelihoods individually.

### Gauge transformation

The effects of natural selection are determined by differences in fitness values, such as the difference between the fitness of the wild type and a mutant. Shifting the fitness values globally by adding a constant, *F* (***g***) ← *F* (***g***) + const., has no effect on fitness differences. In the additive fitness model, it is easy to see that shifting the selection coefficient at any locus by an arbitrary constant *K*_*i*_ does not alter the relative fitness landscape: Σ*i*Σ*a*(*s*_*i*_(*a*) *− K*_*i*_)*δ*_*gi,a*_ = Σ*i s*_*i*_(*g*_*i*_) + const. In other words, the effective fitness parameters can be reduced to (*q −* 1)*L*, and the degrees of freedom that can be arbitrarily adjusted without changing the overall fitness picture are *L* parameters. More systematic arguments under general situations exist and are known as gauge theory in physics and mathematical physics. These concepts have been applied to many genetic sequence-based inference problems^84–86^, with recent reviews for gauge theory in more complex cases^87,88^. In our study, mutation effects of the wild type or TF’s allele serve as reference values; therefore, considering any effects involved with TF’s alleles being zeros is a reasonable choice and makes the inference results more interpretable, as inferred parameters become sparser. To fix the gauge, we employed the following gauge transformation, which is commonly used in statistical inference for genetic sequences^89,90^,

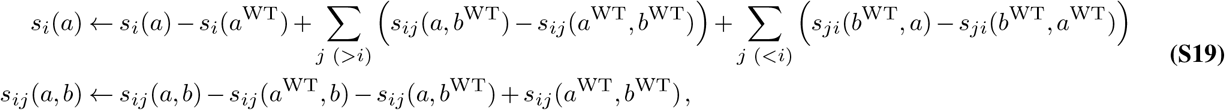

where *a*^WT^, *b*^WT^ are WT (i.e., TF) alleles at locus *i* and *j*, respectively. This choice of gauge ensures *s*_*i*_(*a*^WT^) = *s*_*ij*_(*a*^WT^, *b*) = *s*_*ij*_(*a, b*^WT^) = 0 for all *a, b*.

### Further compression of Ξ

Although the size of the matrix Ξ ∈ ℝ^*D×d*^ is much smaller than the size of the full covariance matrix ∈ ℝ^*D×D*^ with *d* ≪ *D*, still keeping Ξ can be the major bottleneck. An example is HIV-1 CH848 data, where >1200 sequences were collected sequencing more than half of the HIV-1 genome. When we naively compute Ξ storing float variables, that requires more than a terabyte of memory. To further reduce the memory usage, we only consider alleles with nonzero frequency changes Σ_*k*_ |Δ*x*_*i*_(*t*_*k*_)| *>* 0. This modification is straightforwardly implemented in our program.

### Heterogeneous regularization

Instead of applying constant regularization across all parameters, we use a generalized heterogeneous regularization approach with the HCMF method: 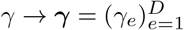. By denoting Λ_***γ***_ as a diagonal matrix with (Λ_***γ***_)_*ef*_ = *γ*_*e*_*δ*_*ef*_, then the optimal fitness parameter becomes *ŝ*= (*C* +Λ_***γ***_)^*−*1^Δ***x***. Consequently, the expression for the efficient expression becomes:

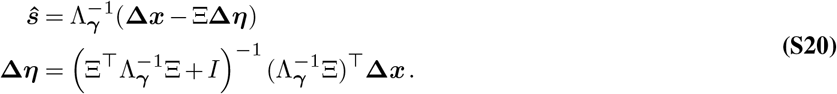

### Theoretical error bars

The posterior distribution is given by the Bayes’ rule,

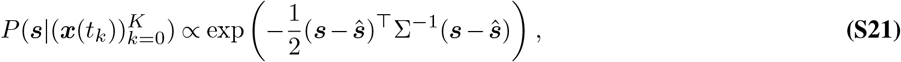

which is a normal distribution for the fitness parameters ***s*** with mean ***ŝ*** and a precision matrix Σ^*−*1^ = (*C*^int^ +*γI*). In a Bayesian inference framework, the uncertainty in the inferred fitness parameters is characterized by (*C*^int^ + *γI*)^*−*1^. More specifically, considering the diagonal of the covariance matrix as the theoretical “error bar,” the standard deviation for *s*_*e*_ can be given by ((*C*^int^ + *γI*)^*−*1^)_*ee*_. Let 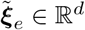 for *e* ∈ {1,…, *D}* be row vectors for Ξ, then by exploiting the structure of the integrated covariance matrix *C*^int^ = ΞΞ^⊤^, one can get

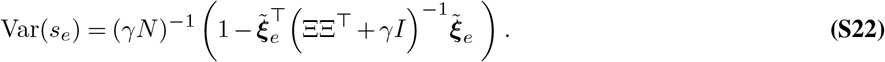

As the inverse in (Eq. (S22)) is easier to obtain, once the inverse is obtained, the variance of each *s*_*e*_ should be straightforwardly obtained.

### Data and code

Data and code used in our analysis is available in the GitHub repository located at https://github.com/bartonlab/paper-hcmf. This repository also contains scripts that can be run to reproduce our figures and analysis.

---

**Supplementary Fig. 1.**
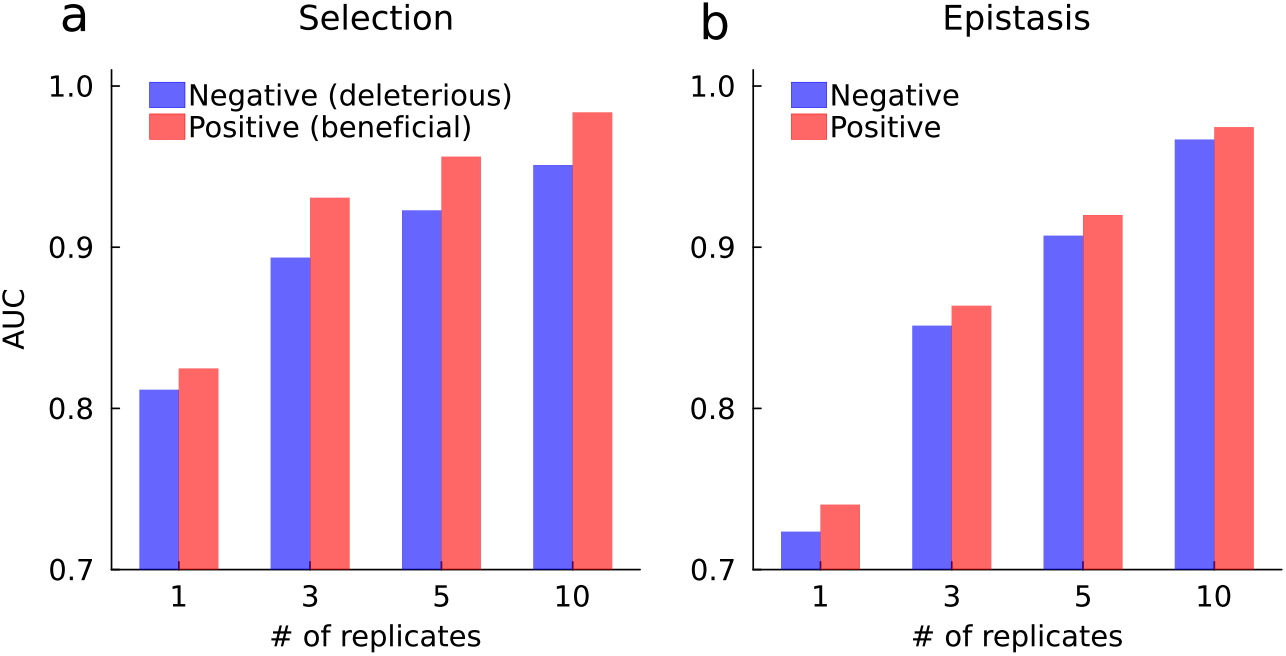
Combining evolutionary replicates improves inference accuracy. **a**, AUC values for identifying selection coefficients as a function of the number of replicates. When using a single trajectory, the AUC values for beneficial and deleterious coefficients are 0.82 and 0.81, respectively. The AUC values for the inferred selection coefficients increase to 0.93 and 0.89, respectively, when combining two replicates. AUC values continue to increase as the number of replicates grows, reaching 0.98 and 0.95 for beneficial and deleterious coefficients with a set of 10 replicates. **b**, AUC values for identifying epistasic interactions from a single replicate are 0.74 for positive and 0.72 for negative epistasis. Similar to the case for selection coefficients, inference accuracy steadily improves with the addition of more replicates.

**Supplementary Fig. 2.**
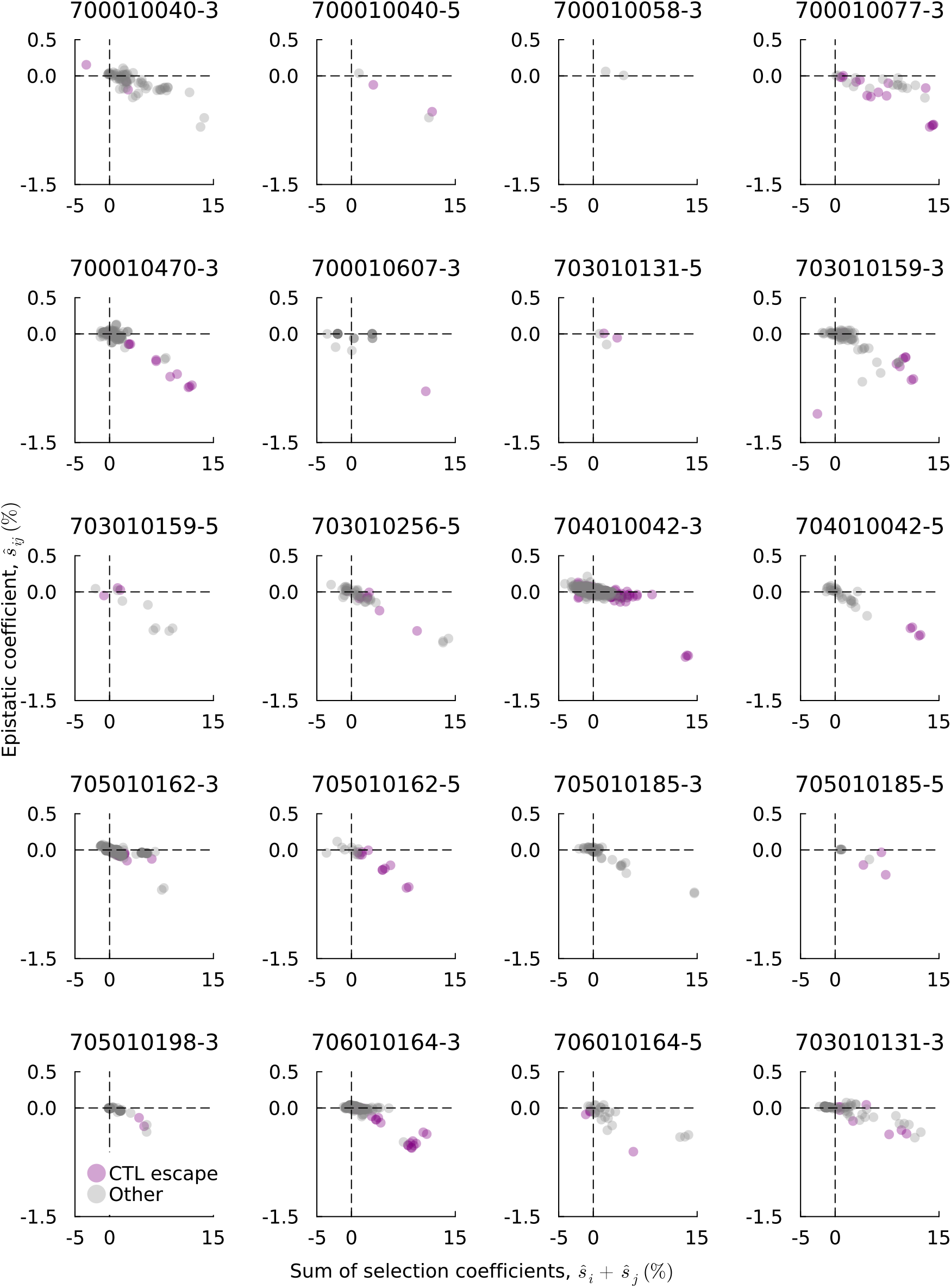
Comparison of inferred epistatic coefficients and the sum of selection coefficients. These figures are analogous to **Fig. 4a** in the main text, but for all individuals and sequencing regions that we analyzed. The tendency of strong anticorrelation between epistasis and the sum of selection coefficients is widely observed across multiple individuals. Relatively strong negative epistatic coefficients are often seen among significantly beneficial mutations, many of which are involved in CTL escape.

**Supplementary Fig. 3.**
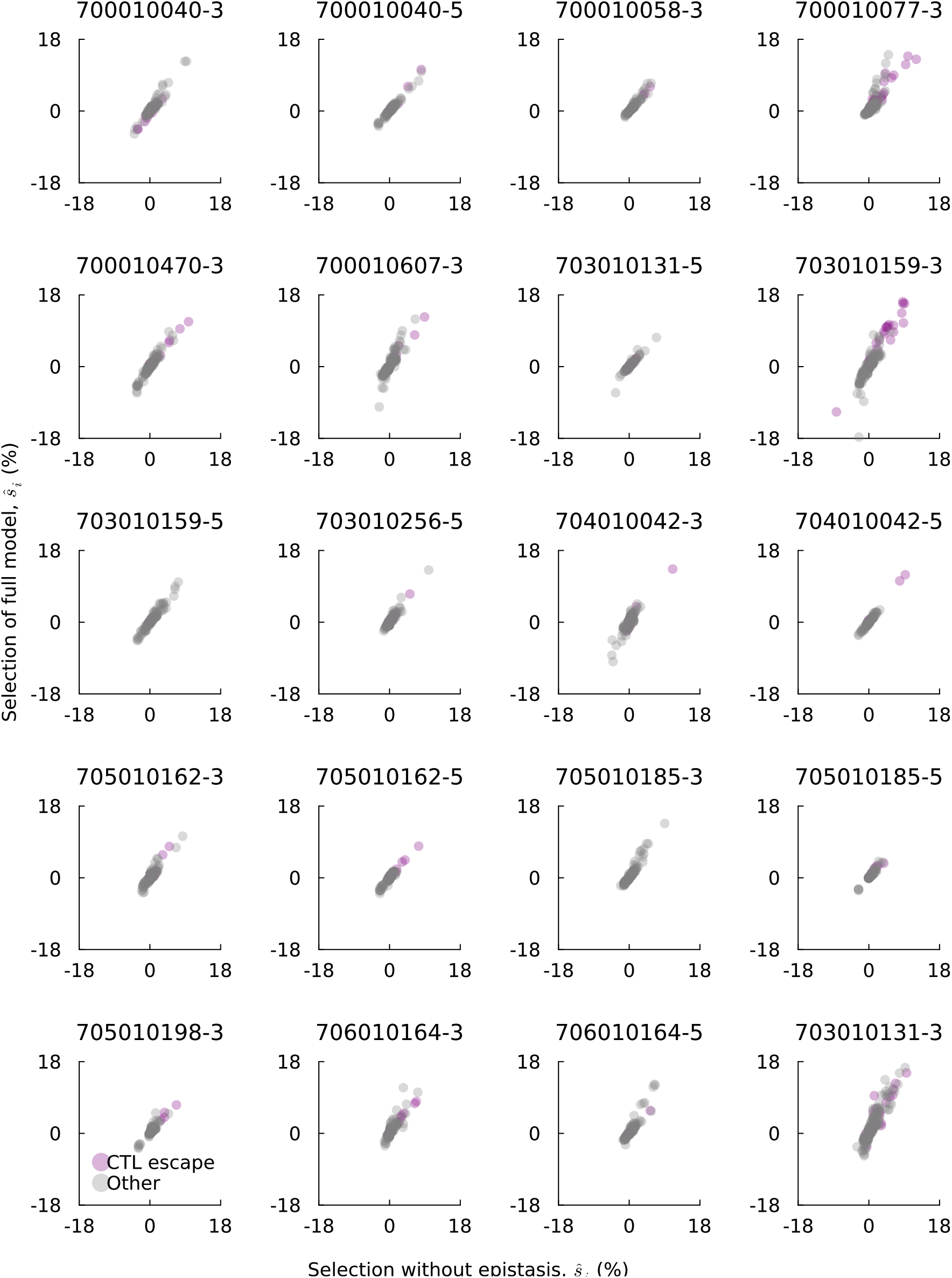
Comparison of inferred selection coefficients in models with and without epistasis. This figure is analogous to **Fig. 5a** in the main text, but across other individuals and sequencing regions that we analyzed. The inferred selection coefficients learned with epistasis are globally consistent with those learned without epistasis. Relatively strong positive selection coefficients are often involved in mutations in CTL epitopes.

**Supplementary Table 1.** Computational time and required memory for inferring epistatic and selection coefficients. The table summarizes the number of polymorphic sites (length *L*), the effective matrix dimension to be inverted (*D*), the number of time points (*K*), as well as the required computational time and memory size for each individual. IDs consist of the patient identifier and sequencing region (5^*′*^ or 3^*′*^ end of the genome), separated by a dash. Computations were performed using a single CPU core with a single thread. For comparison, lengths of around *L* ∼ 200 are already near computational limits even using a high-performance computing system.

| ID | Length, $L$ | Dimension, $D$ | # of time points, $K$ | # of sequences, $N$ | Time (sec) | Memory (GB) |
| --- | --- | --- | --- | --- | --- | --- |
| 700010040-3 | 303 | $1.4 \times 10^5$ | 8 | 82 | 6.2 | 2.6 |
| 700010040-5 | 146 | $3.2 \times 10^4$ | 8 | 74 | 3.5 | 0.7 |
| 700010058-3 | 90 | $1.2 \times 10^4$ | 4 | 25 | 2.6 | 0.3 |
| 700010058-5 | 96 | $1.3 \times 10^5$ | 8 | 52 | 2.4 | 0.4 |
| 700010077-3 | 203 | $6.5 \times 10^4$ | 5 | 44 | 3.7 | 0.8 |
| 700010077-5 | 48 | $3.4 \times 10^3$ | 4 | 32 | 2.3 | 0.3 |
| 700010470-3 | 367 | $2.2 \times 10^5$ | 6 | 113 | 10.8 | 4.8 |
| 700010470-5 | 193 | $5.7 \times 10^4$ | 7 | 104 | 5.9 | 1.4 |
| 700010607-3 | 239 | $8.5 \times 10^4$ | 4 | 73 | 5.2 | 1.5 |
| 700010607-5 | 78 | $9.1 \times 10^3$ | 4 | 76 | 2.3 | 0.4 |
| 703010131-3 | 744 | $8.7 \times 10^4$ | 9 | 114 | 50.8 | 19.4 |
| 703010131-5 | 261 | $9.9 \times 10^4$ | 9 | 76 | 5.0 | 1.9 |
| 703010159-3 | 477 | $3.5 \times 10^5$ | 8 | 98 | 15.8 | 7.0 |
| 703010159-5 | 216 | $7.0 \times 10^4$ | 8 | 93 | 5.5 | 1.6 |
| 703010256-3 | 463 | $3.5 \times 10^5$ | 6 | 99 | 14.7 | 6.6 |
| 703010256-5 | 402 | $2.4 \times 10^5$ | 6 | 110 | 14.1 | 5.5 |
| 704010042-3 | 875 | $1.3 \times 10^6$ | 6 | 93 | 51.1 | 21.6 |
| 704010042-5 | 266 | $1.1 \times 10^5$ | 6 | 85 | 5.1 | 2.1 |
| 705010162-3 | 508 | $4.0 \times 10^5$ | 5 | 69 | 12.6 | 5.7 |
| 705010162-5 | 254 | $9.6 \times 10^4$ | 5 | 60 | 4.9 | 1.4 |
| 705010185-3 | 292 | $1.3 \times 10^5$ | 5 | 97 | 7.3 | 2.7 |
| 705010185-5 | 85 | $1.1 \times 10^4$ | 3 | 49 | 2.3 | 0.4 |
| 705010198-3 | 204 | $6.3 \times 10^4$ | 3 | 48 | 4.1 | 0.9 |
| 705010198-5 | 72 | $7.8 \times 10^3$ | 3 | 47 | 2.2 | 0.3 |
| 706010164-3 | 485 | $3.7 \times 10^5$ | 6 | 102 | 19.1 | 7.4 |
| 706010164-5 | 204 | $6.2 \times 10^4$ | 6 | 98 | 5.1 | 1.5 |
| cap256-3 | 204 | $1.3 \times 10^6$ | 6 | 98 | 5.1 | 1.5 |
| 703010505 | 1,131 | $2.5 \times 10^6$ | 23 | 578 | 319.1 | 215.3 |
| 703010848 | 2,694 | $1.4 \times 10^7$ | 31 | 1,205 | 1,736.0 | 395.5 |

